# Comparative Analysis of Small Secreted Peptide Signaling during Defense Response: Insights from Vascular and Non-Vascular Plants

**DOI:** 10.1101/2024.05.13.593875

**Authors:** Irina Lyapina, Daria Ganaeva, Eugene A. Rogozhin, Ekaterina V. Ryabukhina, Dmitry Yu. Ryazantsev, Vassili Lazarev, Sabina E. Alieva, Anna Mamaeva, Igor Fesenko

## Abstract

Small secreted peptides (SSPs) play an important role in modulating immune responses in all land plants. However, the evolution of stress peptide signaling in different plant phyla remains poorly understood. Here, we compared the expression of SSP genes in the pathogen-induced transcriptomes of vascular and non-vascular plants. We found 13, 19, 15, and 28 SSP families that were differentially expressed during infection in *Physcomitrium patens*, *Zea mays*, *Brassica napus*, and *Solanum tuberosum*, respectively. A comparative study of peptide motifs and predicted three-dimensional structures confirmed the similarity of SSPs across the examined plant species. In both vascular and non-vascular plants, however, only the RALF peptide family was differentially regulated under infection. We also found that EPFL peptides, which are involved in growth and development processes in angiosperms, were differentially regulated in *P. patens* in response to pathogen infection. The search for novel immune-specific peptides revealed a family of PSY-like peptides that are differentially regulated during infection in *P. patens*. The treatment with synthetic EPFL, MEG, PSY, and PSY-like peptides validated their roles in the immune response and growth regulation. Thus, our study showed the complex nature of SSP signaling and shed light on the regulation of SSPs in different plant lineages during infection.

## 1 Introduction

Peptides regulate different processes, including the response to stress conditions (Hou et al., 2021; Lyapina et al., 2019; Olsson et al., 2019). Like animal cytokines (J.-M. Zhang & An, 2007), a group of plant peptides modulating immune response was named "phytocytokines" (Hou et al., 2021). Most of the phytocytokine families were identified in angiosperms. The well-known phytocytokines include PLANT ELICITOR PEPTIDES (PEPs), PAMP-INDUCED PEPTIDES (PIPs), RAPID ALKALINIZATION FACTOR (RALF), PHYTOSULFOKINES (PSKs), PLANT PEPTIDE CONTAINING SULFATED TYROSINE1 (PSY1), SERINE-RICH ENDOGENOUS PEPTIDES (SCOOPs), GOLVENs (GLVs), CLAVATA3/EMBRYO SURROUNDING REGION (CLV3/CLE), CLAVATA3/EMBRYO SURROUNDING REGION (CLV3/CLE), and INFLORESCENCE DEFICIENT IN ABSCISSION (IDA) families (Rzemieniewski & Stegmann, 2022). In addition to the stress response, phytocytokines are shown to be involved in the regulation of plant growth and development (Hou et al., 2021).

Besides phytocytokines, antimicrobial peptides (AMPs) play an important role in plant defense, triggering pathogen cell lysis due to lipid membrane disruption (Moravej et al., 2018). Plant AMPs are small, cysteine-rich proteins (Kulaeva et al., 2020; Tam et al., 2015) that have diverse structures, characteristics, and action mechanisms. The classification of plant AMPs is mostly based on sequence similarity and structures. The most characterized AMPs include thionins, defensins, hevein-like peptides, knottins, hairpin-like peptides, lipid transfer proteins, snakins, and cyclotides (Lima et al., 2022). Antimicrobial peptides were found in most plant taxa.

The phytocytokines, antimicrobial peptides, and plant peptide hormones that control growth and development are often referred to as Small Secreted Peptides (SSPs). The plant SSPs are involved in cell-to-cell communication and share common features: 1) the active peptide sequences are embedded at the C-terminus of 100–250 aa functional or non-functional protein precursors; 2) the protein precursors contain a hydrophobic N-terminal secretion signal; 3) the peptides are released by specific proteases under certain conditions (de Bang et al., 2017; Hu et al., 2021). Nevertheless, precursors of widely recognized phytocytokines, including systemin (SYS) and PEP precursors, do not possess an export signal, which highlights the shortcomings of the SSP classification (Hu et al., 2021; Tavormina et al., 2015). The group of SSPs consists of post-translationally modified (PTM) peptides, cysteine-rich peptides (CRP), and non-Cys-rich and non-PTM peptides. The families of PTM peptides such as PSK, PSY, RGF/GLV/CLEL, CLE, and IDA are widely conserved and were presented in the most recent common ancestor of angiosperms and bryophytes. The cysteine-rich peptides, such as RALF, EPIDERMAL PATTERNING FACTOR (EPF), and EPF-LIKE (EPFL), are shown to be involved in the regulation of many aspects of plant growth and development (Tian et al., 2022). These families were also present in the most recent common ancestor of angiosperms and bryophytes (Furumizu & Shinohara, 2024). The families of SYS and PEP peptides are the most well-known phytocytokines, not belonging to the aforementioned two categories.

During the past decade, thousands of SSPs belonging to more than 70 families have been identified in different plant genomes (Boschiero et al., 2020). In *Triticum aestivum*, 4,981 putative SSPs were identified, among which 1,790 SSPs were grouped into 38 known SSP families (Tian et al., 2022). In *Oryza sativa*, 236 SSPs, including 52 previously unannotated ones, were shown to be involved in the immune response (P. Wang et al., 2020). In *Medicago truncatula*, 4439 genes were predicted to encode SSPs (Boschiero et al., 2020). The bioinformatic search in three bryophyte lineages — mosses, liverworts, and hornworts identified hundreds of SSPs, confirming the important role of peptide signaling in all plant lineages (Lyapina et al., 2021).

The databases of plant SSPs, such as PlantSSP (Ghorbani et al., 2015), OrysSSP (Pan et al., 2013), and MtSSPdb (Boschiero et al., 2020) have been published in recent years. Nevertheless, the catalog of plant SSPs is far from complete, and new peptide families are still being discovered. Recently, a new family of plant phytocytokines, SCREW/CTNIP, was identified in Arabidopsis based on the analysis of stress-induced transcriptomes (Z. Liu et al., 2022; Rhodes et al., 2022). Moreover, plant genomes contain tens of thousands of small open reading frames (smORFs, < 100 aa) that can encode unannotated secreted peptides or be a raw material for the evolution of novel SSPs (Fesenko et al., 2021; Hanada et al., 2013). Such translatable smORFs can be located on transcripts that were annotated as long non-coding RNAs (lncRNAs) (Hsu & Benfey, 2018).

The evolution and role of SSPs in land colonization by plants are currently being discussed (Bowman, 2022; Bowman et al., 2017; Ghorbani et al., 2015). The analysis of phytocytokines in different plant groups showed that most of them are family- or species-specific. That probably reflects their fast evolution, which is driven by arms races between plants and microbes. For example, the SCOOP peptide family is specific to *Brassicaceae* family (Yang et al., 2023). Systemin and hydroxyproline-rich systemin were identified only in *Solanaceae* family (Ryan & Pearce, 2003). A species-specific *Z. mays* immune signaling peptide 1 (Zip1) was found in *Zea mays* (Ziemann et al., 2018). The PEP family is present in many angiosperm families, but their sequences are highly divergent (Lori et al., 2015; Tanaka & Heil, 2021).

Nevertheless, such PTM phytocytokine families as PSK, PSY, RGF/GLV/CLEL, CLE, and IDA were likely present in the most recent common ancestor of angiosperms and bryophytes (Furumizu & Shinohara, 2024). The role of RALF peptides in the immune response was recently demonstrated in both angiosperms and in the moss *P. patens* (Mamaeva et al., 2023). Nevertheless, it is currently unclear whether plant lineages such as bryophytes, lycophytes, ferns, and gymnosperms utilize the same SSP families to modulate immune responses, and how many lineage-specific immune peptide families remain unidentified. Moreover, the understanding of the interactions between several families of phytocytokines in the immune response in diverse plant lineages is currently incomplete.

In this study, we compared the patterns of SSP expression under pathogen invasion in three angiosperm species — Z*ea mays*, *Brassica napus*, and *Solanum tuberosum* — and one bryophyte, the moss *Physcomitrium patens*. Our analysis revealed different patterns of SSP transcription between vascular plants as well as between vascular and non-vascular plants. In addition, our bioinformatic search identified two novel PSY-like peptide families that are specific to bryophytes.

## 2 Materials and Methods

### 2.1 Pre-processing

Raw RNA sequencing data from the following plants under biotic stress were analyzed: the model organism, moss *Physcomitrium patens*, and the angiosperms: *Zea mays*, *Brassica napus*, and *Solanum tuberosum* (accessions: SRP274010, SRP390856, SRP053361, SRP066006, respectively). The following time points were chosen: 4, 8, and 24 hours post inoculation (hpi) for moss *P. patens* infected by *Botrytis cinerea*, 24, 40, 60 hpi for *Z. mays* infected by *Colletotrichum graminicola;* 24, 48, 96 hpi for *B. napus* infected by *Sclerotinia sclerotiorum*; 12, 24, 72 hpi for *S. tuberosum* infected by *Pectobacterium carotovorum*.

FastQC (Andrews, 2010) was used for quality control, and Trimmomatic (Bolger et al., 2014) was used to clip adapters and nucleotides of low quality. Genomes and genome annotations were accessed via Phytozome (https://phytozome-next.jgi.doe.gov/, Goodstein et al., 2012) and Ensembl Plants (https://plants.ensembl.org/). *P. patens* v3.3, *Z. mays* Ref_Gen_V4, *B. napus* AST_PRJEB5043_v1, and *S. tuberosum* v6.1 genome versions were selected.

### 2.3 Assembly and ORF prediction

Genomes were indexed using hisat2-build, reads were mapped to the reference genome using HISAT2 (Kim et al., 2015). Samtools (H. Li et al., 2009) was used for collecting mapping statistics.

*De novo* transcript assembly was done using Stringtie (Pertea et al., 2015), Stringtie -e mode was applied for quantification. Prediction of open reading frames (ORFs) from AUG start codons for assembled transcripts, along with further automatic translation to amino acid sequences longer than 30 aa, was performed using orfipy (Singh & Wurtele, 2021). The longest ORF per transcript was selected, considering that the longest ORF represents actual coding sequence.

### 2.4 Functional annotation

Interproscan (Quevillon et al., 2005, version 5.66-98.0) was used to predict protein domains. Analysis of homology with known families of small secreted peptides (SSPs) was done using the SSP prediction tool (https://mtsspdb.zhaolab.org/prediction/, Boschiero et al., 2020). Plant long non-coding RNAs were accessed via PLncDB (https://www.tobaccodb.org/plncdb/).

### 2.5 DE analysis and filtration

Differential gene expression analysis was done using the edgeR package (Robinson et al., 2010). Transcripts with 1 > logFC < -1 and FDR < 0.05 were considered differentially expressed.

ORFs predicted as known SSPs by the SSP prediction tool were assigned to the corresponding SSP family based on homology to SSPs from other plants. The search for potential novel candidates of SSPs was conducted in ORFs shorter than 200 aa, corresponding to differentially regulated transcripts, with predicted signal sequences, no transmembrane domains, no known protein domains, and no homology to known SSPs from other plants.

### 2.6 Sequence alignment and structure prediction

Multiple sequence alignments (MSAs) were created using the high precision L-INS-I algorithm of MAFFT v7 (Katoh & Standley, 2013). Alignments were visualized using JalView v2.11.2.2, Clustalx color (Waterhouse et al., 2009).

Prediction of three-dimensional structures was done using AlphaFold2 (Jumper et al., 2021). Structural comparison was performed using TM align (Y. Zhang & Skolnick, 2005). Such metrics as RMSD and TM-score were considered. Structural search against PDB and AlphaFold database (AFDB) was performed using Foldseek (van Kempen et al., 2024).

### 2.7 Search for peptide homologs

A search for paralogs of novel immune peptide candidates was performed using BLASTP against all major ORFs assigned to each transcript. Hits with E-value < 0.00001 were considered significant. The search for orthologs and paralogs for novel candidates in other databases was done using NCBI BLASTP against the non-redundant protein sequences database, organism: *Viridiplantae.* Sequences with no matches or with matches to unannotated proteins (“putative”, “unknown”, “uncharacterized”, etc) were selected. TBLASTN against 1000 plant transcriptomes (1KP, https://db.cngb.org/datamart/plant/DATApla4/) was run to search for homologs of novel candidates among translated nucleotide sequences of other plants published in 1KP. The following default parameters were customized: a maximum of 250 target sequences and a word size of 3. Hits with E-value < 0.01 were considered significant.

### 2.8 Cloning and recombinant protein expression

The pET32b+ vector (“Novagen”, USA) was used for recombinant plasmids construction. When the obtained recombinant plasmid is expressed, a hybrid protein is created that contains thioredoxin, polyhistidine domain, enteropeptidase cleavage site (DDDDR) and the peptide sequences. Previously, an assembly of synthetic genes encoding MEG and EPFL peptides was carried out by PCR method. This procedure was made according to Shevchuk et al. (2004): the long oligonucleotides (Supplementary Table 1) (100 ng of each) and all required components were added to PCR mixture (the Phusion HS II DNA polymerase was used). The PCR amplification was carried out using the following program: 94°C – 60 s; 94°C – 20 s, 60°C – 20 s, 72°C – 20 s – 10 cycles; 72°C – 60 s. Then, 2.5 µl of the obtained mixture was added to a novel amplification mixture containing flanking oligonucleotides for the corresponding peptide (0.5 µM of each), and the amplification was carried out according to the same program (20 cycles). The PCR product was purified from the agarose gel and cloned into the pET32b+ vector using *HihdIII/KpnI* restriction endonucleases. The obtained construction was transformed into *E. coli* strain Origami. The induction and purification of the recombinant protein were carried out under native conditions according to the QIAexpressionist manual (“Qiagen”, Germany). For the purpose of fusion protein purification from bacterial cells 1000 ml LB growth medium was inoculated with 25 ml of overnight *E. coli* culture. The induction was carried out for 4 h in the presence of 1 mM IPTG, then the cells were centrifuged, and the precipitate was resuspended in 50 ml of buffer A (50 mM NaH_2_PO_4_, 300 mM NaCl, 10 mM imidazole, pH 8.0). The *E. coli* cells disruption was realized for 30 min on ice with the addition of lysozyme (1 mg/ml) followed by sonication. The purified cell lysate was applied to Ni-NTA agarose column (10 ml) pre-equilibrated with buffer A, then the column was washed with buffer A containing 20 mM imidazole. The fusion protein was eluted with buffer B (50 mM NaH_2_PO_4_, 300 mM NaCl, 250 mM imidazole, pH 8.0). The purity and size of recombinant hybrid proteins were evaluated by SDS-PAGE electrophoresis.

### 2.9 Peptide chemical synthesis

*P.patens* PSY1, PSYL1, and PSYL2 peptides (Table 1) were chemically synthesized at the genetic engineering laboratory of the FRCC PCM (Moscow, Russia). The peptides were synthesized using solid-phase synthesis on Rink resin. The synthesis was carried out with microwave irradiation of the reaction mixture, according to the Fmoc-strategy. The purity of the lyophilized peptides was > 95%, their molecular weight was confirmed by mass spectrometric analysis. The synthesized peptides were dissolved in sterile water and stored at − 80°C.

**Table 1.**
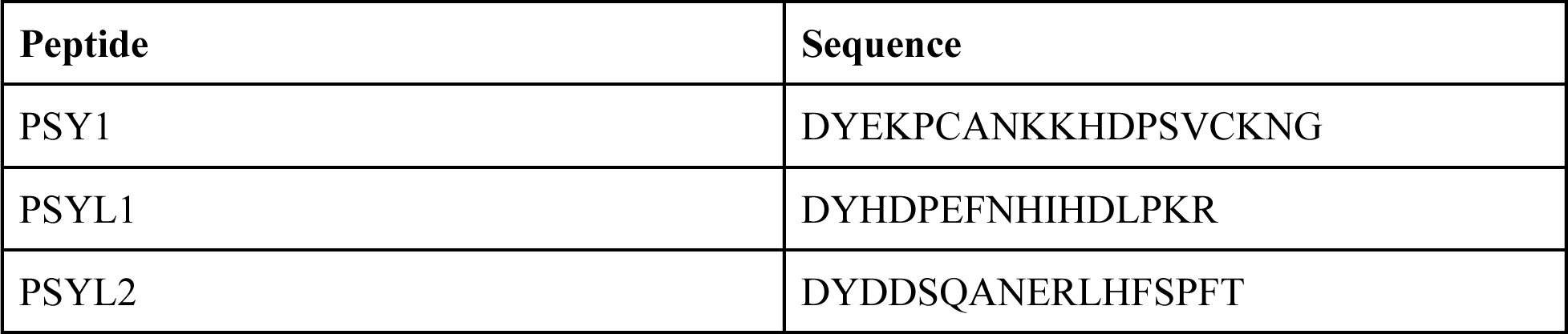
Sequences of synthesized peptides.

### 2.10 Plant growth

Protonemata of *Physcomitrium patens* subsp. patens Gransden 2004 (Freiburg, Germany) were grown in 100 ml liquid Knop medium containing 500 mg/l ammonium tartrate for ROS accumulation analysis or in 600 µl solid Knop medium with 450 µl sterile water containing a corresponding peptide, ammonium tartrate and glucose for cell length analysis, under white light with a photon flux of 61 μmol/m2 s under a 16-hour photoperiod at 24°C. Seven-day-old protonema was used for analysis.

### 2.11 ROS detection and cell length analysis

For intracellular ROS detection, we used the fluorescent dye 2ʹ,7ʹ-Dichlorofluorescin Diacetate (DCFH-DA; Sigma-Aldrich, USA). *P. patens* protonemata were treated with 5 µM synthesized PpEPFL or PpMEG for 10 min. Samples were incubated with 10 µM DCFH-DA for 15 min. The detection was performed on the fluorescent microscope Axio Imager M2 (Zeiss) with an AxioCam 506 mono digital camera and filter units. The No. 44 filter (λex BP 475 nm/40 nm; λem BP 530 nm/50 nm) was used for DCFH-DA fluorescence detection. Data on the fluorescence intensity were obtained from the related Zeiss software Zen.

For cell length analysis, protonema was grown with 1 nM - 1 µM of each peptide for 7 days. Cell walls were stained with 10 µg/ml propidium iodide for 5 min. Filter unit 20 Rhodamin (λex BP 546/12 nm; λem BP 575–640 nm) was used.

### 2.12 Statistics

Statistical analysis and visualization were made in Python v. 3.7.4 (Van Rossum and Drake, 1995) using modules scipy 1.5.2 (Virtanen et al., 2020), seaborn 0.11.1 (Waskom, 2021), numpy 1.20.1, pandas 1.2.3 (McKinney, 2012). For determining statistically significant differences two- or more-way analysis of variance (ANOVA), followed by Tukey’s honestly significant difference (HSD) test, or Mann-Whitney U test were applied. Differences were considered significant at a P-value ˂ 0.05.

## 3 Results

### 3.1 Identification of SSPs in pathogen-induced transcriptomes

In order to analyze the expression patterns of known and previously unannotated SSPs during immune response, we re-analyzed four published transcriptomes of phylogenetically distant species: the moss *Physcomitrium patens* (Physcomitrella), *Zea mays*, *Brassica napus*, and *Solanum tuberosum* (Kwenda et al., 2016; Reboledo et al., 2021; D. Wang et al., 2022; Xia et al., 2023; Figure 1A). These transcriptomes were selected based on the following criteria: 1) the plants were infected with either necrotrophic or hemibiotrophic pathogens; 2) the quality of raw data (see Methods); and 3) RNA-seq analysis was performed at different time points after pathogen inoculation.

**Figure 1.**
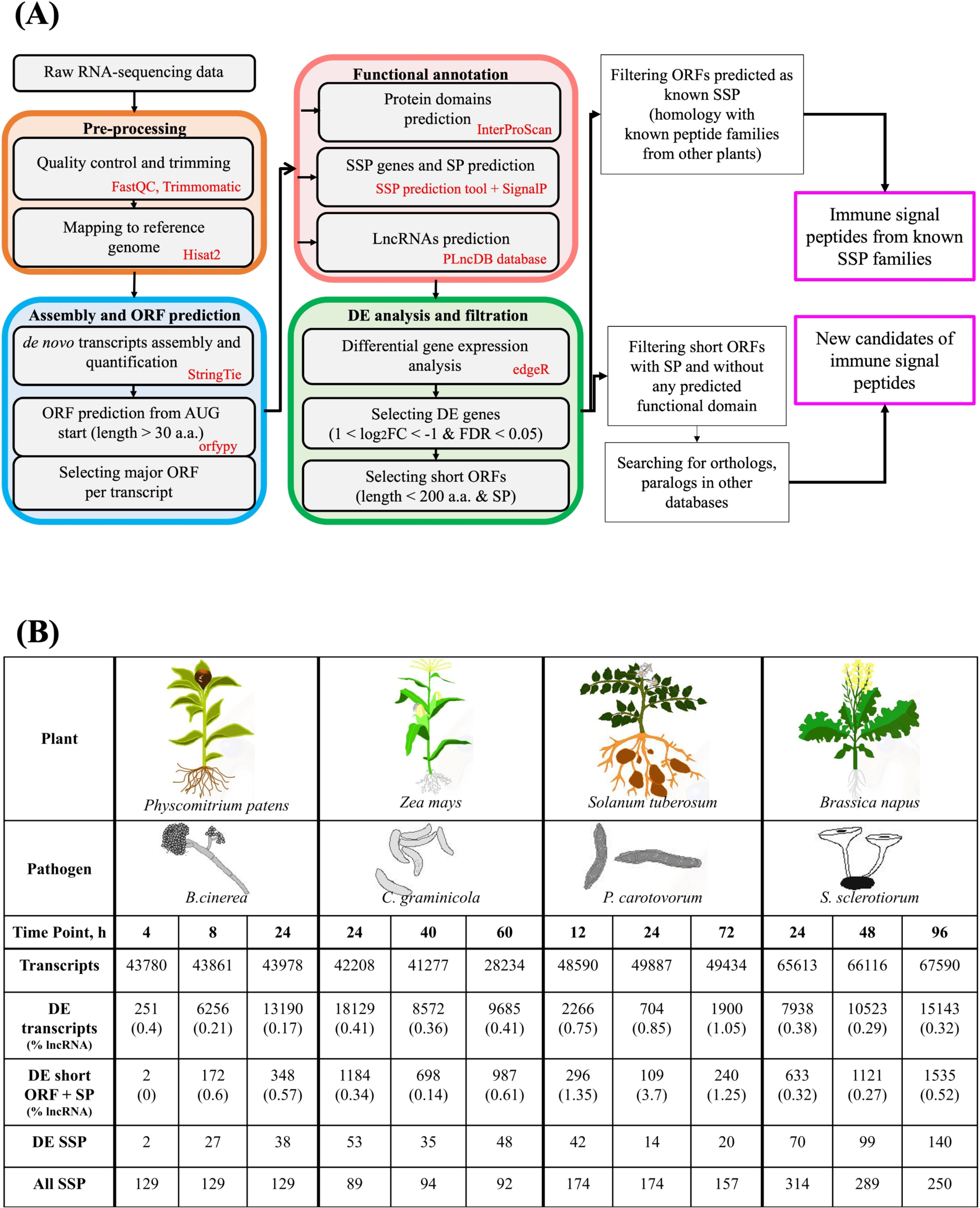
**(A) -** General scheme of work pipeline; **(B) -** transcripts assembly statistics.

The RNA-seq reads were remapped to the corresponding genomes with the following alignment rate: 91% for *P. patens*; 96% for *Z. mays*; 83% for *B. napus*; and 85% for *S. tuberosum*. Overall, we identified from 28,234 to 67,590 transcripts at different hpi (hours post infection) across all selected species (Figure 1B). The following number of transcripts were assigned to annotated genes: 23,843 for *P. patens*; 15,269 for *S. tuberosum*; 15,138 for *Z. mays* and 19,875 for *B. napus*. To annotate all assembled transcripts, open reading frames (ORFs) larger than 30 amino acids (aa) that start with AUG codons were predicted using the orfipy tool (Singh & Wurtele, 2021). For annotation, the major ORF was selected for each transcript, resulting in approximately 45,000 predicted proteins in *P. patens* and *S. tuberosum*; approximately 40,000 and 60,500 predicted proteins in *Z. mays* and *B. napus*, respectively.

The predicted proteins were then annotated using the InterProScan tool (Quevillon et al., 2005). This resulted in assigning to known protein domains about 57% of the assembled transcripts in *P. patens* and *B. napus*, 53% in *S. tuberosum* and 70% in *Z. mays*. Among the most represented domains, such as chloroplast- or ribosome-related domains, a significant portion of the predicted ORFs belonged to uncharacterized domains or were unannotated (Supplementary Table 2).

We next used the online SSP prediction tool (https://mtsspdb.zhaolab.org/prediction/, Boschiero et al., 2020) to determine the transcripts encoding putative precursors of known SSP families (Figure 1B; Supplementary Table 3). Overall, 129, 92, 168, and 284 transcripts encoding potential SSP precursors were identified in the assembled transcriptomes of *P. patens*, *Z. mays*, *S. tuberosum*, and *B. napus*, respectively (Figure 1B; Supplementary Table 3).

For each species, we next identified differentially expressed (DE) transcripts (-1 ≤ log2 fold change ≥ 1, *P*adj < 0.05) at each time point (Figure 1B; Supplementary Table 2). In *P. patens* and *Z. mays*, the highest number of DE transcripts was found at 24 hpi - 30% and 43% of all transcripts, respectively (Figure 1B; Supplementary Table 2). For *S. tuberosum* the largest number of DE transcripts was found at 12 hpi (4.7% of overall transcripts), and for *B. napus* - at 96 hpi (22.4% of overall transcripts) (Figure 1B; Supplementary Table 2).

We additionally identified lncRNAs in assembled transcriptomes that could potentially encode for unannotated peptide precursors (Hsu & Benfey, 2018). To estimate the proportion of lncRNAs in differentially regulated transcripts, we selected the assembled transcripts that 1) do not code for annotated proteins; 2) do not assign to known protein domains and SSP families; 3) contain smORFs below 100 codons; and 4) do not overlap known genes on the same strand. We annotated such transcripts using the lncRNA database PLncDB (https://www.tobaccodb.org/plncdb/). At each time point, lncRNA transcripts accounted for approximately 0.17–1.05% of differentially regulated transcripts in each species, with *S. tuberosum* having a higher percentage and *P. patens* having a lower one (Figure 1B; Supplementary Table 2). We suggest that it can be the result of both biological and technical reasons.

For further analysis, we selected 13, 19, 15, and 28 DE SSP families from *P. patens*, *Z. mays*, *S. tuberosum,* and *B. napus*, respectively (22, 45, 25 and 103 DE transcripts, respectively; Figure 1B; Supplementary Table 3). Moreover, we additionally selected DE unannotated proteins (≤ 200 aa), containing N-terminal signals (Figure 1B; Supplementary Table 2). This set of DE proteins was used for the identification of novel, previously unannotated peptide families. Some of them were located on predicted long noncoding RNAs (Figure 1B; Supplementary Table 2).

### 3.2 Transcriptional patterns of SSP genes in different plant lineages

We next analyzed the transcriptional patterns of different groups of SSPs in response to phytopathogen infection. The first group consists of selected known phytocytokines. The second group included "peptide hormones," which have been demonstrated to control growth and development but not stress responses. The third group consists of known families of AMPs.

#### 3.2.1 The transcriptional regulation of phytocytokines

The most common differentially regulated phytocytokines in all examined species were from conserved phytocytokine families such as PSK, CLE, PSY, and RALF. The transcripts that encode RALF peptides were identified in all examined species at three time points. The members of this family were differentially expressed under infection in *P. patens*, *B. napus* and *Z. mays*. In *P. patens*, the PpRALF1 (Pp3c3_15280) gene was significantly upregulated under infection (Table 2; Supplementary Table 3). Some members of the RALF family were significantly downregulated under infection in *B. napus* and *Z. mays* (e.g., CDX78035 in *B. napus* and Zm00001d033941_T001 in *Z. mays*; Table 2).

**Table 2.**
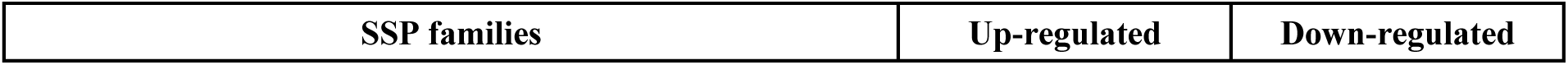
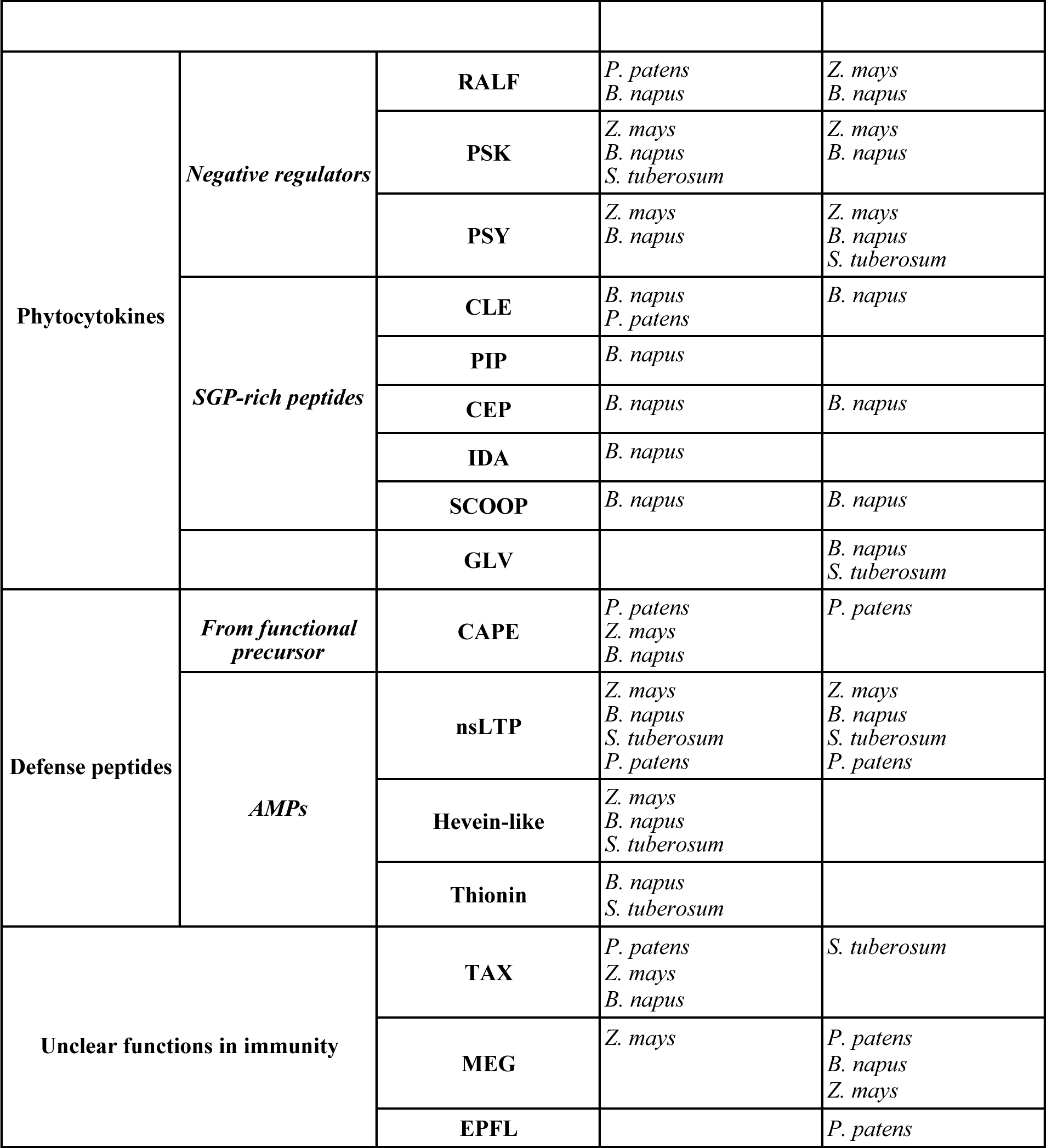
Groups of differentially expressed gene families predicted as known SSPs involved in immunity.

Previously, PSK peptides were shown to regulate the immune response to biotrophic and necrotrophic pathogens in an antagonistic manner in Arabidopsis (Y. Li et al., 2024). The genes belonging to the PSK family were differentially regulated in vascular plants but not in *P. patens*. In Z. *mays*, PHYTOSULFOKINES 6-RELATED gene (Zm00001d023385_T001) was upregulated during infection. In addition, another phytosulfokine gene (Zm00001d023384_T001) was up- and downregulated at different hpi, and Zm00001d048006_T001 was downregulated. In *B. napus*, we identified 22 upregulated transcripts of PSK genes, and two genes, CDX68551 and CDY36518, were downregulated. One PSK gene (Soltu.DM.02G027440.2) was upregulated in *S. tuberosum* (Table 2; Supplementary Table 3).

The PSY family was recently shown to play a role in regulating the trade-off between plant growth and stress response in vascular plants (Ogawa-Ohnishi et al., 2022). PSY receptors (PSYRs) activate the expression of various genes encoding stress response transcription factors upon depletion of the ligands (Ogawa-Ohnishi et al., 2022). We found that PSY transcripts were significantly downregulated in all vascular plants under infection: MSTRG.18847.1 in *S. tuberosum*; CDY71288, CDX80347, and MSTRG.42712.1 in *B. napus*; 14 transcripts in *Z. mays* (Table 2; Supplementary Table 3). Some PSY genes in *Z. mays* and *B. napus* were also upregulated: Zm00001d038523_T001, Zm00001d020443_T001, MSTRG.18595.2 and MSTRG.17209.2 in *Z. mays*; CDY64688, CDY35659, CDY69338 and MSTRG.43937.2 in *B. napus* (Table 2; Supplementary Table 3). However, our analysis did not confirm the expression of the predicted PSY gene in P. *patens*.

The analysis of phytocytokines specific to vascular plants identified PIP (7 upregulated transcripts), CEP (12 upregulated transcripts), IDA (CDY08563), and SCOOP (three upregulated and one downregulated) peptides in *B. napus* (Table 2; Supplementary Table 3). Transcripts of IDA peptide precursors were detected in *S. tuberosum* plants, but their transcription did not show significant changes under infection.

Transcripts of PEP precursors (PROPEPs) were not detected in our analysis across the examined plant species, likely due to their low transcription levels.

Thus, our analysis showed that only the RALF peptide family was differentially regulated under infection in vascular and non-vascular plants.

#### 3.2.2 The transcriptional patterns of known peptide growth hormones

The number of SSP families that were shown to regulate only growth and development processes is limited. For example, cysteine-rich EPIDERMAL PATTERNING FACTOR (EPF) and EPF-LIKE (EPFL) peptide families play a role in regulating plant growth and development (Y. He et al., 2023; Xiong et al., 2022). Recently, the EPFL family was found to be significantly downregulated upon *F. oxisporum* infection and elicitor treatments in tomato (Slezina et al., 2021). Our analysis revealed transcripts encoding members of the EPFL family in all angiosperms, although their transcription levels remained unchanged during infection. However, we found that some members of the EPFL family were significantly downregulated in the moss *P. patens*. According to our analysis, 14 transcripts belonging to the EPFL peptide family were identified in *P. patens*. From them, nine transcripts were significantly downregulated (six annotated genes: Pp3c6_12270, Pp3c23_5720, Pp3c24_9860, Pp3c23_11350, Pp3c1_26030, Pp3c6_27020; Table 2; Supplementary Table 3). The function of this peptide family in bryophytes is currently unknown. Thus, these results suggest that moss EPFL peptides may be involved in the regulation of the immune response, as has been recently shown in tomato (Slezina et al., 2021).

The genes encoding members of the cysteine-rich TAX (TAXIMIN) family peptide were significantly upregulated in *Z. mays* (four transcripts, three isoforms of Zm00001d008923_T001 and one not annotated MSTRG.18593.1), *B. napus* (seven transcripts, two annotated: CDY31955 and CDY19064), *P. patens* (Pp3c4_17790) and downregulated in *S. tuberosum* (MSTRG.21268.1; Table 2; Supplementary Table 3). This family was suggested to play a role in nutrient-status-related signaling (de Bang et al., 2017).

Another peptide family of cysteine-rich peptides, MEG (Maternally Expressed Gene), was previously shown to be downregulated in tomato plants under infection (Slezina et al., 2021). We found that members of this family were significantly downregulated in *P. patens* (two isoforms of Pp3c5_6640) and *B. napus* (MSTRG.22520.1) and some members were significantly up- and down-regulated in *Z. mays* (three up-regulated transcripts: two isoforms of Zm00001d005037_T001 and Zm00001d052375_T002; two down-regulated transcripts: Zm00001d052375_T002 and Zm00001d042365_T001; Table 2; Supplementary Table 3).

TPD 1 (TAPETUM DETERMINANT 1) is a secreted cysteine-rich small protein that was shown to control anther cell differentiation in Arabidopsis (Huang et al., 2016). We identified two TPD transcripts in the moss *P. patens* (STRG.20746.1, STRG.20746.2) and three of the five transcripts encoded TPD-like peptides in *Z. mays* (two of them already annotated (Zm00001d049823_T001, Zm00001d006677_T001) were significantly up-regulated (Supplementary Table 3).

Thus, our analysis shows that cysteine-rich peptides, which were not considered to play a role in regulating immune responses, were actively regulated during infection in all plant lineages. Our findings show that the widely conserved RALF and EPFL peptide families may play a role in the immune response in both vascular and non-vascular plants.

#### 3.2.3 The transcriptional regulation of antimicrobial SSPs

We identified several different SSPs classified as antimicrobial and protease inhibitors in the analyzed species, such as Gibberellic acid stimulated in Arabidopsis (GASA) from Snakin family, ARACIN, Potato type II proteinase inhibitor, SubIn and others. The most known antimicrobial peptides, hevein-like and thionins, were identified in the transcriptomes of all examined species (Supplementary Table 3). However, hevein-like transcripts were significantly upregulated during infection only in vascular plants (Table 2; Supplementary Table 3). Thionins were significantly upregulated in *B. napus* and *S. tuberosum*, but we did not identify differential expression in other species.

The non-specific lipid transfer proteins (nsLTPs) are a plant-specific superfamily of cysteine-rich AMPs (Santos-Silva et al., 2023). The transcripts of nsLTP genes were identified in all analyzed plant species and were the most highly represented group of transcripts, both up- and down-regulated (Supplementary Table 3). Our findings showed that nsLTPs have similar patterns of regulation in response to pathogen attack in vascular and non-vascular plants (Table 2).

### 3.3 Comparative analysis of SSP motifs

We next compared the peptide motifs of bryophyte and angiosperm SSPs differentially regulated in the transcriptomes of the examined species. The SSPs were selected based on the following criteria: 1) differentially expressed in *P. patens* and in at least one angiosperm species; or 2) differentially expressed in *P. patens*. We used only functional peptide sequences for this analysis (Figure 2A; Supplementary Figure 1).

**Figure 2.**
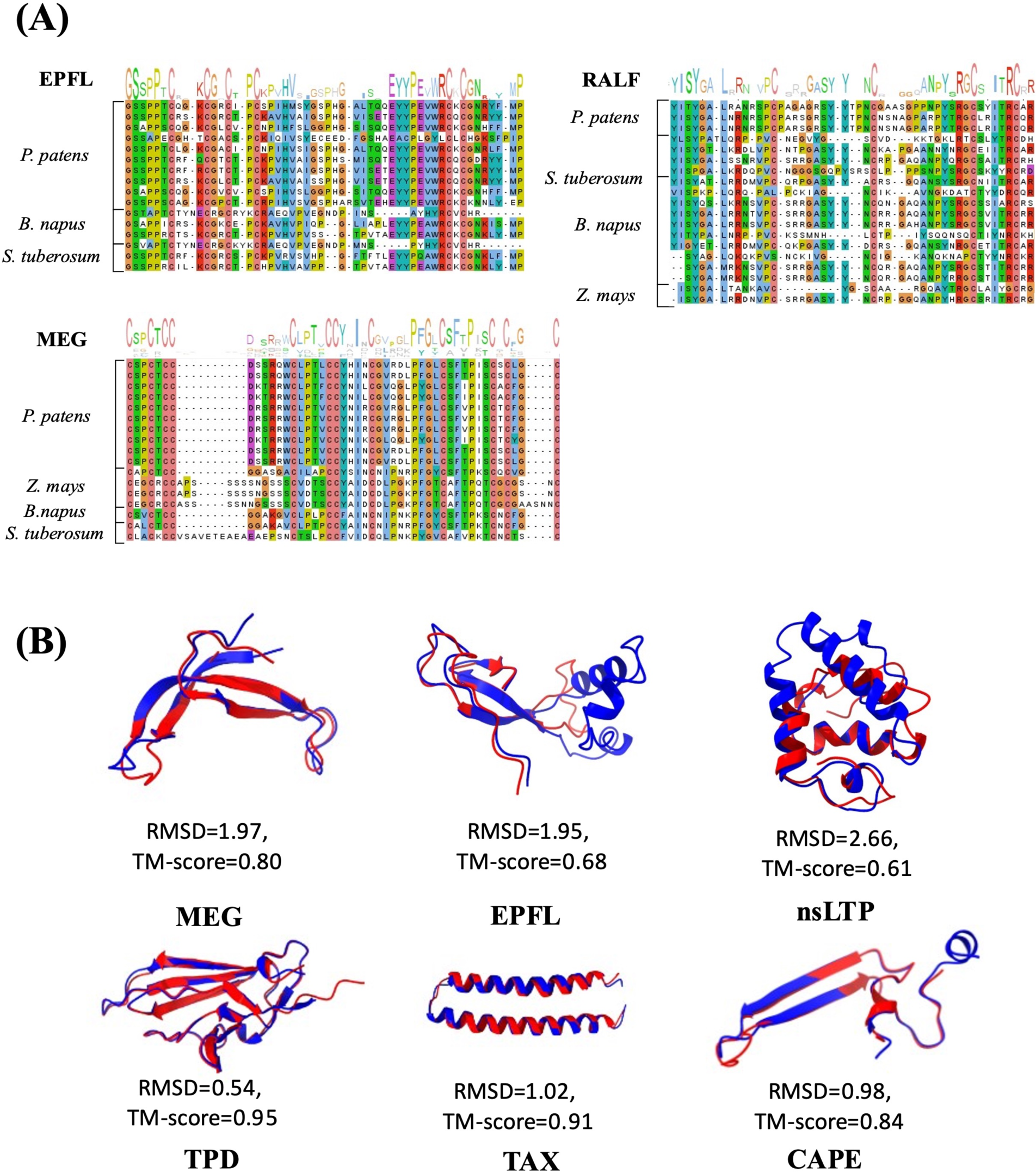
**(A) -** Sequence similarity analysis for peptide families in *Physcomitrium patens* and angiosperms; (**B) -** Structural comparison of angiosperm and *Physcomitrium patens* peptide predicted structures.

The EPFL peptides identified in *P. patens*, *S. tuberosum*, and *B. napus* revealed a typical cysteine pattern as well as a conserved GS region (Figure 2A, Supplementary Figure 1A), which is similar to the previous studies (Takata et al., 2013; Xiong et al., 2022).

The YISY motif, which is shown to play a role in receptor binding (R. Zhang et al., 2023), was identified in most RALF peptides from *P. patens*, *S. tuberosum,* and *B. napus*. *Z. mays* peptides contain an ISY motif (Figure 2A, Supplementary Figure 1B). The RGC region was found in all studied plants, suggesting that RALF peptides share known motifs in all examined species (R. Zhang et al., 2023).

Previously, MEG peptides with 8 and 12 cysteine residues were identified in angiosperms (Gutiérrez-Marcos et al., 2004; Slezina et al., 2021). We found that MEG peptides from *P. patens*,*Z. mays*, *S. tuberosum,* and *B. napus* contain 12 conserved cysteine residues, including two CC regions (Figure 2A, Supplementary Figure 1C). In contrast to the cysteine patterns, the MEG amino acid sequences showed significant variation among the examined sequences.

Thus, our analysis showed that EPFL peptides are the most conserved among all the plants analyzed. The largest difference in peptide motifs was observed in MEG peptides, including the number of cysteine residues and overall amino acid composition.

### 3.4 Structure-based comparison of the identified SSPs

#### 3.4.1 The prediction of peptide structures using AlphaFold2

The accuracy of sequence-based methods for predicting peptide structures has been demonstrated in recent years (McDonald et al., 2023). Moreover, current methods for structure comparison allow the detection of distant orthologs (Monzon et al., 2022). Therefore, we next compared predicted structures of EPFL, RALF, CLE, MEG, nsLTP, CAPE, TPD, and TAX peptides from different species. For this, 135 peptide structures were predicted by AlphaFold2 (Jumper et al., 2021), and structures with an average pLDDT > 70 were selected for further analysis. Peptide structures of *P. patens* were compared with those of angiosperms. Structural alignment was performed using the TM-align tool, and metrics including sequence identity (Seq_Id), Root Mean Square Deviation (RMSD), and TM-score were calculated (Supplementary Table 4). Because clear secondary structures were not predicted for RALF and CLE peptide, they were excluded from further analysis. Overall, 21 peptide structures of MEG, EPFL, CAPE, TPD, TAX, and nsLTP peptides with the highest average pLDDT per family for each plant species were selected for structural comparison (Supplementary Table 4, Supplementary Materials).

The predicted structures of the MEG peptides consist of multiple beta-sheets. The highest similarity to *P. patens* MEG structures was observed in the case of *Z. mays* MEG (TM-score 0.80; Figure 2B), which is in line with our motif comparison. The lowest structural similarity was found between *P. patens* and *S. tuberosum* MEG structures (TM-score 0.44, Supplementary Table 4). The MEG structure of *Z. mays* contains an additional loop composed of alpha-helices, which is absent in the predicted MEG structure of *P. patens*.

The structure of EPFL peptides comprises two antiparallel beta-sheets connected by a loop. The structures of EPFL peptides from *Z. mays* and *P. patens* showed the highest similarity (TM-score 0.68; Figure 2A) compared to other analyzed plants (TM-scores 0.55 for *B. napus* and *S. tuberosum;* Supplementary Table 4).

The predicted structure of non-specific lipid transport proteins (nsLTPs) contains four alpha-helices. *B. napus* nsLTP peptide structure revealed the highest similarity to *P. patens* ndLTP structures (Figure 2B). TM-scores for *Z. mays* and *S. tuberosum* structures compared to *P. patens* were 0.50 and 0.56, respectively (Supplementary Table 4).

The predicted structures of the TPD and TAX peptides showed high structural similarity across all examined species (Supplementary Table 4). The highest similarity between the predicted structures of *P. patens* and *B. napus* peptides was observed (Figure 2B).

The predicted structures of the CAPE peptides contain two beta-sheets and an alpha-helix. The highest similarity to *P. patens* CAPE was observed between P. *patens* and *B. napus* structures (Figure 2B).

Therefore, the cysteine-rich peptides (MEG, EPFL) from *P. patens* showed the highest similarity to *Z. mays* homologs. The structures of CAPE, nsLTP, TAX, and TPD peptides from *P. patens* were the most similar to the predicted structures from *B. napus*. The TPD peptide has the most conserved structure among *P. patens, Z. mays,* and *B. napus* species.

#### 3.4.2 The search for structural homologs of identified SSPs

Then, we searched for structural homologs of *P. patens* peptide families MEG, EPFL, CAPE, nsLTP, TAX, and TPD against AlphaFold Protein Structure Database (https://alphafold.ebi.ac.uk) and PDB (https://www.rcsb.org) databases by Foldseek (van Kempen et al., 2024).

The analysis of MEG peptides yielded three significant hits (TM-score > 0.5, Probability > 0.8) against the AlphaFold Protein Structure Database (AFDB) with CRP7-Cysteine-rich family protein from *Zea mays* (Supplementary Figure 2).

For EPFL peptides, structural similarity to ∼70 EPFL proteins from *Oryza sativa, Glycine max, Arabidopsis thaliana, Nicotiana tabacum, etc.* was identified. Hits to structures from the PDB database were also obtained, such as EPFL and plant receptor ERL1-TMM in complex with EPF1 from *Arabidopsis thaliana* (Accessions in PDB database: 2liy_A, 5xkj_F, 5xjo_E; Supplementary Figure 2). We also found hits to the structures of nsLTP, TPD, TAX, and CAPE peptides in the AFDB and PDB databases (Supplementary Figure 2).

Thus, our analysis revealed high structural similarity among MEG, EPFL, CAPE, nsLTP, TAX, and TPD peptides from different plant taxonomic groups. For CAPE and EPFL precursors, hits to PDB structures were observed. Therefore, our results confirmed that searches by Foldseek (van Kempen et al., 2024) can be used to find peptide homologs at the structural level.

### 3.5 Identification of novel families of SSPs in the moss *P. patens*

Novel peptide families have emerged during the evolution of land plants (Furumizu & Shinohara, 2024). However, the identification of taxon-specific peptides and novel peptide families is a challenging task. Moreover, lineage-specific peptide families have been identified only in angiosperms to date. Therefore, we next sought to search for novel SSPs involved in the immune response in the non-vascular plant, *P. patens,* using our custom pipeline.

At first, we selected 356 ORFs below 200 aa located on DE transcripts that contained a predicted N-terminus signal sequence but did not contain a transmembrane domain or known peptide motif (Figure 1). We additionally filtered out “candidate” SSPs that have protein annotation or have hits in the NCBI non-redundant protein sequence database (BLASTP, E-value < 0.01, organism: *Viridiplantae*). We next searched for possible orthologs of 58 “candidate” SSPs in the 1000 plant transcriptome (1KP) database (https://db.cngb.org/datamart/plant/DATApla4/) using TBLASTN (E-value < 0.01). Then, we manually checked the presence of possible peptide motifs based on multiple sequence alignments (MSAs).

Based on the analysis of possible paralogs in the *P. patens* genome, we identified two families of putative novel SSPs in *P. patens*. The consensus sequences of the identified motifs are presented in Figure 3A (Supplementary Figure 3). Family 1 comprises three *P. patens* genes (*Pp3c2_4250*, *Pp3c14_17750*, and *Pp3c21_20040*) that have not been functionally annotated yet. One of these genes, *Pp3c21_20040*, was differentially regulated upon *B. cinerea* infection (Figure 3A, Supplementary Figure 3A). The members of Family 1 are cysteine-rich and comprise 8 conservative cysteine residues as well as conserved “NATH” and “SCGF” motifs (Figure 3A). The TBLASTN search in the 1KP database identified possible orthologs in 49 species, including mosses, liverworts, and lycophytes (Figure 3A, Supplementary Figure 4). The median length of predicted ORFs was 119 aa.

**Figure 3.**
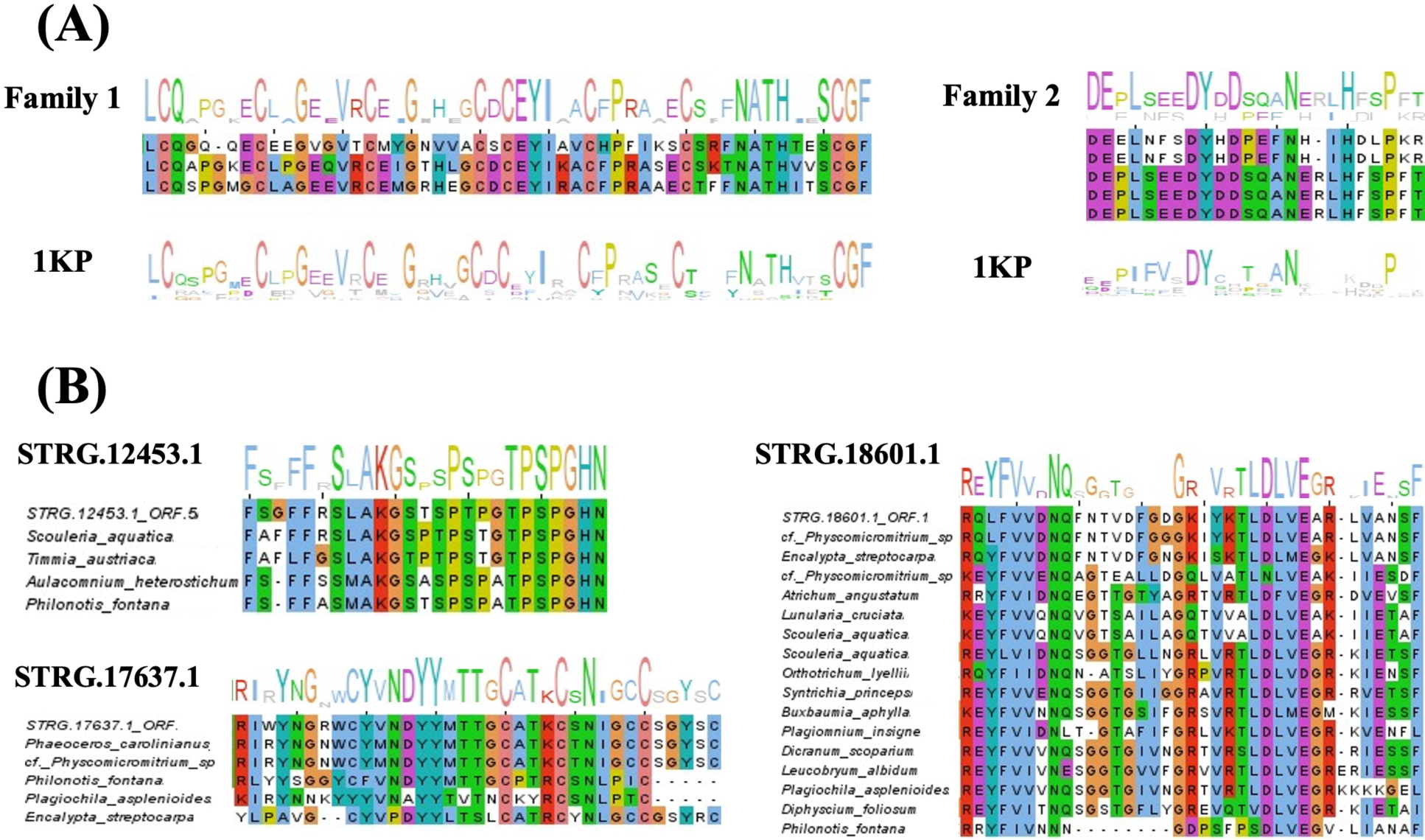
(**A) -** Alignment logos of novel candidate families found in *P. patens* (**Family 1:** STRG.19963.1_ORF.1; STRG.1695.1_ORF.2; STRG.14124.1_ORF.4; alignment logo with 1KP; **Family 2:** STRG.4116.1_ORF.1; STRG.4116.2_ORF.1; STRG.4476.2_ORF.1; STRG.4476.3_ORF.1; STRG.4473.1_ORF.1; alignment logo with 1KP); (**B) -** Alignment logos of novel peptide candidates corresponding to unique *P. patens* ORFs with 1KP (STRG.12453.1_ORF.5; STRG.18601.1_ORF.1; STRG.17637.1_ORF.1).

The putative peptide precursors of Family 2 shared two similar peptide motifs: DY*D***N**H**P and DY*D***N***H**P (Figure 3A, Supplementary Figure 3B). These motifs are similar to the known PSY peptide motif DY*(D)(P)**N**H*P (Furumizu et al., 2021), but differ in the position of conserved residues H and P. Previously, one *P. patens* gene (Pp3c8_3200) was identified as a precursor of PSY peptide (Furumizu et al., 2021; Tost et al., 2021). Two predicted protein precursors from Family 2, STRG.4116.1_ORF.1 (117 aa) and STRG.4116.2_ORF.1 (91aa), shared the DY*D***N**H**P motif at the C-end and came from the *Pp3c3_33940V3.3* gene that was erroneously annotated as alternative splicing factor SRp20/9G8 in Phytozome 13. The STRG.4116.1 transcript was differentially upregulated upon infection with *B. cinerea*. The other three predicted precursors contain the motif DY*D***N***H**P. Two of them, STRG.4476.2_ORF.1 (114 aa) and STRG.4476.3_ORF.1 (118 aa), belonged to the gene *Pp3c4_6140*. The assembled transcript STRG.4473.1 intersects the *P. patens* gene *Pp3c4_6040* and encodes 118 aa precursor STRG.4473.1_ORF.1. The TBLASTN analysis of the corresponding peptide sequences against 1000 Plant transcriptomes identified 20 possible orthologs of Family 2 in mosses and one liverwort species (Supplementary Figure 5). The median length of predicted ORFs was 110 aa which is similar to known PSY genes. Thus, we identified a novel family of immune-responsive PSY-like (PSYL) peptides that are encoded by three *P. patens* loci.

Because SSPs can be presented by only one copy in the genomes of non-vascular plants, we additionally analyzed putative SSP precursors without possible paralogs in *P. patens*. We searched for their possible homologs against transcripts from the 1KP database. The resulting alignments of the three most interesting up-regulated peptide candidates are presented in Figure 3B. Full alignments can be found in Supplementary Figure 6. These putative peptide precursors contain motifs that are conserved among different plant species. In the case of STRG.12453.1_ORF.5 (140aa), P***TPSPGHN motif is observed, which is similar to CLE and IDA peptide motifs (Furumizu et al., 2021). This candidate was found in four moss species (Figure 3B, Supplementary Figure 6A). Candidate STRG.18601.1_ORF.1 (115aa) contains a conserved LDLVE motif, which is observed in 13 other species, including mosses and liverworts (Figure 3A, Supplementary Figure 6B). Finally, *P. patens* candidate STRG.17637.1_ORF.1 (121aa) is cysteine-rich and, notably, contains a conserved DYY region. Hits to four other species were found, including moss, liverwort, and hornwort plants (Figure 3B, Supplementary Figure 6C). Further study is needed to explore the functions of these SSP candidates.

Thus, we propose an approach to identify novel, not annotated, plant peptide families. This approach focuses on the identification of putative SSP precursors (based on length, N-terminal signal sequence, no TM-domain) combined with analysis of possible peptide motifs. Overall, we identified 5 potential candidates for novel signaling peptides in *P. patens*. Two of them comprise families based on sequence similarity, and some of them are unique ORFs predicted in *P. patens.* Nevertheless, all potential novel candidates are found in other plant species belonging to non-vascular plants. For the two candidates, distant homology to known peptide families (PSY, IDA, and CLE) is suspected.

### 3.6 Experimental validation of potential immune peptides

We next synthesized and assessed the biological activity of the moss PSY, PSYL1, PSYL2, PpEPFL, and PpMEG peptides. At first, we used reactive oxygen species (ROS) accumulation as a marker of the immune response. Treatment with 5 µM of synthetic PpPSYL2 and PpEPFL induced a significant accumulation of ROS molecules in *P. patens* cells compared to the mock (Mann-Whitney U test: P-value < 0.0005; Figure 4A). In contrast, treatment with syntenic PpPSY1 and PpMEG resulted in a significant decrease in the accumulation of ROS in moss cells, which may indicate their role as negative regulators of the immune response (Mann-Whitney U test: P-value < 0.0005; Figure 4A). We did not observe any significant difference in the ROS accumulation between the mock and PSYL1 peptide treatments (Figure 4A).

**Figure 4.**
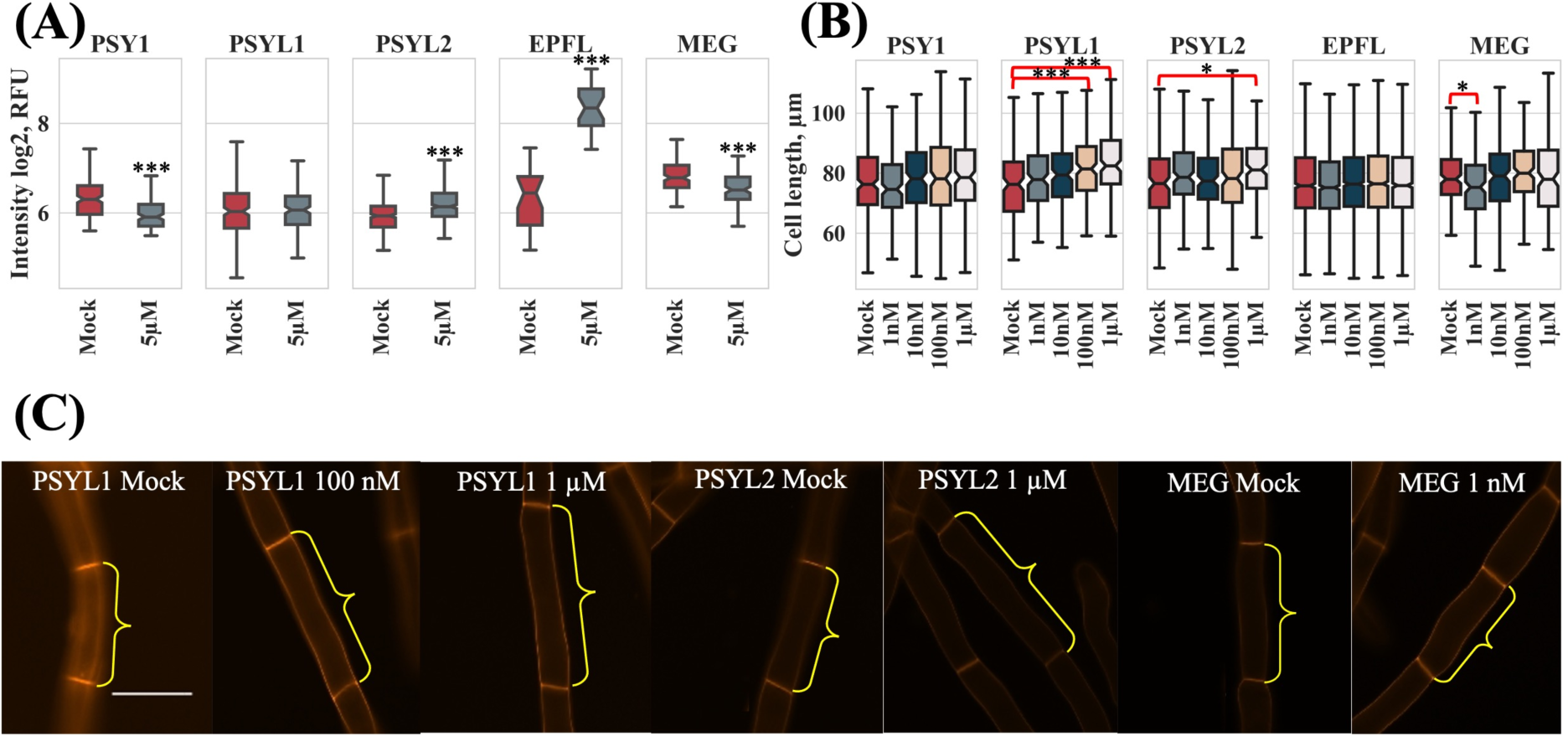
(**A) -** Analysis of the accumulation of intracellular ROS molecules using a fluorescent dye DCFH-DA after treatment with 5 µM PpPSY1, PpPSYL1, PpPAYL2, PpEPFL and PpMEG in the protonema of moss *P. patens* (Mann-Whitney U test: ***P-value < 0.001). Control samples (C) were treated with water. **(B) -** Measurement of the length of protonema cells grown in a semi-solid medium containing PpPSY1, PpPSYL1, PpPSYL2, PpMEG, PpEPFL (Tukey’s test: ***P-value < 0.001; *P-value < 0.05). (**C) -** Microscope images showing moss *P. patens* protonemata cells stained with propidium iodide. Bar = 50 µm. All experiments were performed in at least three replicates.

We also explored the effects of peptide treatment on *P. patens* growth. The moss protonemata were grown on a semi-solid medium supplemented with either a corresponding peptide or a mock treatment for one week. A significant difference in cell length was found between mock and treatment with PSYL1 (100 nM and 1 µM; Tukey’s test: P-value < 0.001), PSYL2 (1 µM), and MEG (1 nM) peptides (Tukey’s test: P-value < 0.05; Figure 4B and C).

Thus, our experiments confirmed the biological activity of differentially regulated SSPs, including previously unknown PSY-like peptides.

## 4 Discussion

Recent studies showed that peptides derived from protein precursors play an important role in plant immunity, modulating the defense response. The plant cell-surface receptors that recognize these peptide ligands contain Leucine-Rich Repeat (LRR) or Malectin ectodomains and have an ancient origin (Ngou et al., 2024). Contrary to immune receptors, the origin and evolution of immune peptides are still poorly understood (Furumizu et al., 2021). One of the reasons is that, due to rapid changes, they appear to quickly enter the "twilight zone", where sequence similarity is no longer detectable. Nevertheless, several databases that include plant Small Secreted Peptides (SSPs) were recently published (Boschiero et al., 2020; Ghorbani et al., 2015; Pan et al., 2013). Sequence similarity, however, does not necessarily mean functional similarity across different plant phyla. In this study, we compared pathogen-induced transcriptomes in vascular and non-vascular plants to understand how SSP families from different plant lineages respond to infection. Along with conserved SSPs associated with immune responses that were found in the transcriptomes of both vascular and non-vascular plants, we observed species-specific regulation of certain SSP families.

Our analysis shows that cysteine-rich peptides were actively regulated during infection in all plant lineages. The EPFL, RALF, TAX, and MEG were differentially regulated in *P. patens* and angiosperm species. The widely conserved family of EPIDERMAL PATTERNING FACTOR (EPF) peptides is involved in regulating plant growth or development in vascular and non-vascular land plants (Caine et al., 2016; Y. He et al., 2023). The nine EPFL genes that are upregulated in the developing sporophyte have been identified in previous bioinformatics studies of the *P. patens* genome (Caine et al., 2016; Lyapina et al., 2021; Takata et al., 2013), but their functions remain unknown. However, the role of EPF and EPF-LIKE (EPFL) peptides as positive or negative regulators of immune response has not been shown to date. Nevertheless, several genes of the EPFL family were shown to be significantly downregulated upon pathogen infection in tomato (Slezina et al., 2021). In this study, we found that the transcriptional level of six EPFL genes was significantly downregulated in *P. patens*. This result can be explained by a specific response to pathogens or by the inhibition of growth processes under stress conditions.

Our treatment of moss protonemata with synthetic EPFL peptide resulted in an increase in intracellular reactive oxygen species (ROS) levels, suggesting the activation of downstream signaling. In Arabidopsis, the receptor-like kinase (RLK) ERECTA and its two closely related homologs, ERECTA-LIKE 1 (ERL1) and ERL2 are responsible for perceiving the EPF/EPFL family peptides (Y. He et al., 2023; Tameshige et al., 2017). In the genome of *P. patens,* six potential orthologues of the ERECTA genes were found (Villagarcia et al., 2012). Thus, the EPFL family might be a target for future work on understanding the evolution of phytocytokines in land plants.

The RALF peptides were shown to be involved in immune response across different plant lineages (Y.-H. He et al., 2023; Mamaeva et al., 2023; Slezina et al., 2021; Wolf & Höfte, 2014). We found both up-regulated and down-regulated RALF genes in pathogen-induced transcriptomes, which is consistent with previous results showing different roles of different members of RALFs in modulating plant immunity (Y.-H. He et al., 2023; L. Liu et al., 2024; Mamaeva et al., 2023; Stegmann et al., 2017).

The poorly characterized cysteine-rich peptides, such as TAX and MEG, were differentially regulated in both vascular and non-vascular plant species. However, how TAX peptides are involved in the immune response is currently unknown. Previously, tomato MEGs were found to be downregulated in response to infection (Slezina et al., 2021). This is similar to our findings and suggests the conservative role of this SSP family in vascular and non-vascular land plants. Nevertheless, the role of MEG peptides in plant growth and defense is poorly understood.

In addition to common approaches for the identification of SSPs based on sequence similarity (Boschiero et al., 2020), we developed a new pipeline to identify novel lineage-specific secreted peptides. Our pipeline resulted in the identification of two putative SSP families in *P. patens*. One of these families (Family 2) consists of PSY-like peptides, which are similar to the known phytocytokine PSY family in angiosperms. The PSY peptide family was present in the most recent common ancestor of vascular plants and bryophytes (Furumizu & Shinohara, 2024). Previously, one gene potentially encoding the PSY peptide and another encoding the cognate PSYR receptor were identified in *P. patens* (Furumizu et al., 2021; Furumizu & Aalen, 2023). Thus, we significantly expanded the family of PSY-like peptides in bryophytes. The identification of previously unknown PSY-like peptides in bryophyte genomes could expand our understanding of the peptide evolution in land plants.

In conclusion, our comprehensive analysis of small secreted peptides in the pathogen-induced transcriptomes of four plant species showed the conservation of cysteine-rich peptides across different plant lineages. We found high motif and structural conservation of these peptides in vascular and non-vascular plants. In addition, we identified previously uncharacterized putative SSP families in bryophytes. One of these bryophyte-specific families shows similarity to previously characterized PSY peptides. Thus, our study provides a new avenue to explore the evolution of known and previously uncharacterized peptide families.

## Author contributions

IF supervised the study. IL, DG conducted all experiments and performed all bioinformatic analyses. AM performed ROS measurement and cell length analysis. EAR, DYR and EVR synthesized recombinant moss *P. patens* peptides. VL and SEA synthesized moss *P. patens* PSY family peptides. IL, DG, and IF wrote the manuscript. All authors contributed to the article and approved the submitted version.

## Acknowledgment

We thank the Center for Precision Genome Editing and Genetic Technologies for Biomedicine, Lopukhin Federal Research and Clinical Center of Physical-Chemical Medicine of Federal Medical Biological Agency for the expertise and guidance in genetic engineering.

## Data availability statement

The data that support the findings were derived from the NCBI Sequence Reads Archive (SRA) available in the public domain with the following accession numbers: SRP274010, SRP390856, SRP053361, SRP066006.

